# Deformable Mirror based Optimal PSF Engineering for 3D Super-resolution Imaging

**DOI:** 10.1101/2022.05.09.491071

**Authors:** Shuang Fu, Mengfan Li, Lulu Zhou, Yingchuan He, Xin Liu, Xiang Hao, Yiming Li

**Affiliations:** Department of Biomedical Engineering, Southern University of Science and Technology, Shenzhen 518055, China; State Key Laboratory of Modern Optical Instrumentation, College of Optical Science and Technology, Zhejiang University, Hangzhou 310027, China

## Abstract

Point spread function (PSF) engineering is an important technique to encode the properties (e.g., 3D positions, color, and orientation) of single molecule in the shape of the PSF, often with the help of a programmable phase modulator. Deformable mirror (DM) is currently the most widely used phase modulator for fluorescence detection as it shows negligible photon loss. However, it relies on careful calibration for precise wavefront control. Therefore, design of an optimal PSF not only relies on the theoretical calculation of maximum information content, but also the physical behavior of the phase modulator, which is often ignored during the optimization process. Here, we developed a framework of PSF engineering which could generate a device specific optimal PSF for 3D super-resolution imaging using DM. We used our method to generate two types of PSFs with depths of field comparable to the widely used astigmatism and Tetrapod PSFs, respectively. We demonstrated the superior performance of the DM specific optimal PSF over the conventional astigmatism and Tetrapod PSF both theoretically and experimentally.

---

Single molecule localization microscopy (SMLM) has successfully overcome the diffraction limit, making it possible for researchers to resolve biological structures with nanoscale resolution[1]. Over the past ~15 years, tremendous efforts have been made to extend this technology from 2D to more dimensions, such as axial position, color information, fluorescence polarization, motion diffusion, and lifetime[2]. Among these developments, precise 3D imaging is probably the most important one. A variety of methods for 3D super-resolution imaging have been devised[3], and PSF engineering is the most used method to obtain the 3D information. Through clever design of PSF, the 3D information of an emitter can be precisely extracted from its shape in the 2D images, even optimized for different imaging range[4].

The simplest and most widely-used PSF engineering method is to insert a cylindrical lens to create an astigmatism PSF with its ellipticity depending on the *z* position within a limited axial range (~1μm)[5]. To extend 3D imaging to a larger axial range, engineered PSFs with more complex pupil functions are normally needed. There are two main ways to generate these engineered PSFs. One is to fabricate specific transmission phase masks for different types of PSFs. Although the requirement of the precision for fabrication of phase mask has been lowered by using liquid immersion recently[6], phase masks lack the flexibility for various biological applications and system aberration correction. A more general alternative way is to employ a programmable phase modulator in the Fourier plane of the optical system. Adaptive optics devices such as spatial light modulator (SLM) and DM are commercially available for this purpose. SLM has the advantage of large number of pixels. Therefore, it can output a complicated phase even with sharp phase jumps (*i.e*., DH-PSF[7]). However, only one polarization of light can be modulated by SLM, halving the precious fluorescence photon budget in SMLM experiments. DM shows negligible photon loss and is very suitable for phase modulation in fluorescence imaging. However, it has limited number of actuators, making it challenge to accurately output the designed phase pattern.

To design optimal PSFs for different imaging conditions, a variety of algorithms have been proposed. First approaches optimized the Zernike coefficients of the pupil function by minimizing the 3D Cramér-Rao lower bound (CRLB) [8][9]. A family of “Tetrapod” PSFs were found to achieve the best theoretical precision within predefined axial range[4]. Recently, deep-learning based methods were also employed to optimize PSFs for dense 3D imaging[10]. By jointly optimizing the PSF generation and localization convolutional neuron network, the parameterized phase mask which yields superior reconstruction for high-density 3D localization could be obtained. Furthermore, heuristic PSFs with analytical pupil functions were also introduced which achieved near optimal performance compared to the Tetrapod PSFs[11]. However, none of these algorithms were designed by taking account for the physical response of the phase modulator used.

Here, we used DM as the phase modulator which is most photon efficient and convenient for fluoresce PSF engineering comparing to the other phase modulation methods. However, calibration of the DM is often a prerequisite that cannot be neglected for precise wavefront control[12][13]. In this work, we carefully calibrated the experimental DM influence function of each actuator by using an interferometer. Instead of optimizing the coefficients of Zernike polynomials, we directly used the influence function of each actuator as basis function of the pupil function optimized. We call the PSF obtained by this way as DM-optimized PSF (DMO PSF). The feasibility of the DMO PSF is verified by high-quality 3D SMLM images, which are reconstructed using either the spline fitting method [14] or deep learning network [15].

As shown in **Fig. 1a**, we equipped our SMLM system with a DM (DM140A-35-P01, Boston Micromachines) placing in the Fourier plane of the microscope, which allowed us to modulate the pupil function with its 140 actuators (**Fig. S1**). For the practical DM calibration, we follow the method in Ref. [16], which utilizes a Twyman-Green interferometer to measure the surface deformation of the DM. Here, we only calibrated the influence function of each actuator instead of a set of Zernike modes (**Supplementary Note 1**). Comparing to the calibrated Zernike modes, the influence functions can offer more accurate wavefront design as they represent original physical activation of the actuators and avoid the approximation errors which often happens during the computation of the Zernike modes.

**Fig. 1.**
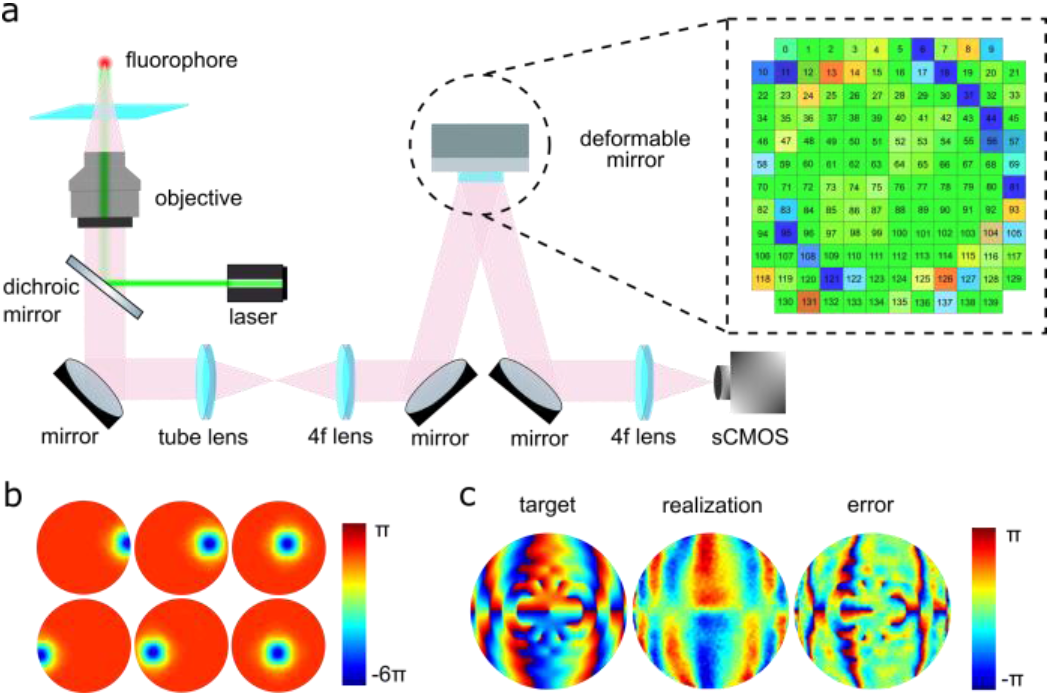
(a) Schematic of the PSF optimization using a deformable mirror, inset is the control panel of a DM with 140 actuators. Color indicates the relative voltage applied to each actuator. (b) The example influence functions of the DM. (c) Projection of the pupil function of DH-PSF on DM.

**Fig. 1b** shows the calibrated influence functions of a DM for a few represented actuators. As shown in **Fig. 1c**, we decomposed the phase of a DH-PSF using DM influence functions. The residual error is quite large, limiting the practical usage of some complex PSFs with DM. Therefore, it is important to take account for the response of the DM actuators when designing a PSF engineered by DM. It is worth to note that the effect of an influence function spreads over neighboring actuators (**Fig. 1b**). Therefore, the total 140 influence functions were used for the optimization process, although the aperture of the objective often does not fill the whole active area of the DM.

Since CRLB corresponds to the limit of attainable precision of single molecule localization, we optimize the DM-engineered phase *ψ_DM_* with respect to CRLB:

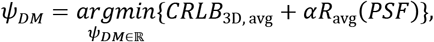

where 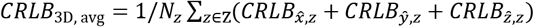 is the averaged 3D CRLB over *N_z_* discrete *z* positions in a predefined axial range Z. In this work, we used *CRLB*_3D, avg_ with *z* step of 100 nm for 1.2 μm axial range PSF and 200 nm for 6 μm axial range PSF.

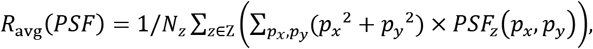

where (*p_x_*, *p_y_*) represents pixel coordinates with the center of PSF model as zero. *R*_avg_(*PSF*) term is used to confine the spatial extent of the PSF to reduce overlap. *α* is a hyper-parameter, which we set as 0 and 30 for the 1.2 μm and 6 μm PSF optimization, respectively. It is worth to note that one can also use different weights for CRLB xy and z to give different importance between lateral and axial localization precision. To implement the optimization, we used the explicit gradient derivation (**Supplementary Note 2** and **3**) and the interior-point method of the *fmincon* function in Matlab.

Considering that searching the DM control signal in a high dimensional parameter space (140 dimensions for our DM) is time-consuming and prone to fall into a local minimum, we first searched for a solution in a relatively low dimensional space using all 21 tertiary Zernike polynomials (2<n+ļmļ ≤ 8, where *n* is the radial order and *m* is the angular frequency) as the basis functions. This Zernike based pupil function was then decomposed using DM influence functions to obtain the realistic pupil function by DM. We then continue to optimize the pupil function in the space of the calibrated DM influence functions. We optimized with different starting points. Nevertheless, the DMO PSF normally showed better *CRLB*_3D, avg_ than that of the PSF generated by Zernike based optimal pupil function on DM (~6%-23%, **Fig. S2**). This is because the projection of the Zernike based pupil function to the DM would introduce approximation errors.

Most of 3D SMLM applications utilize astigmatism for 3D imaging as it is easy to be implemented and data analysis is also quite simple even compatible with elliptical Gaussian PSF[4] **(Fig. 2a**). However, its 3D imaging capability is not optimal. Therefore, we try to use our approach to optimize a PSF with a similar imaging depth as a normal astigmatism PSF (~1.2 μm). The resulting pupil function and theoretical PSF are shown in **Fig. 2b**. We call this PSF as DMO Saddle-point PSF. Compared to the astigmatism PSF, DMO Saddle-point PSF shows more concentrated intensity distribution than the astigmatism PSF when defocus, thus improving the overall *CRLB*_3D, avg_. The localization precision of astigmatism PSF only shows slightly better resolution near the focus (+-140 nm). However, its performance decreases dramatically when the imaging plane is bit away from the focus, as the localization precision is more than 3 times worse between focus and ±600 nm axial positions (**Fig. 2c** and **d**). In comparison, DMO Saddle-point PSF maintains a relatively uniformed resolution over the axial range being optimized. This property is quite important for the quantitative analysis of biological structures over a relatively large imaging range. Furthermore, the localization precision of x and y for an astigmatism PSF is asymmetry about the focus which could potentially distort the reconstructed image even in 2D. The localization precision of DMO Saddle-point PSF showed preferable symmetry resolution along the x and y direction at all axial positions.

**Fig. 2.**
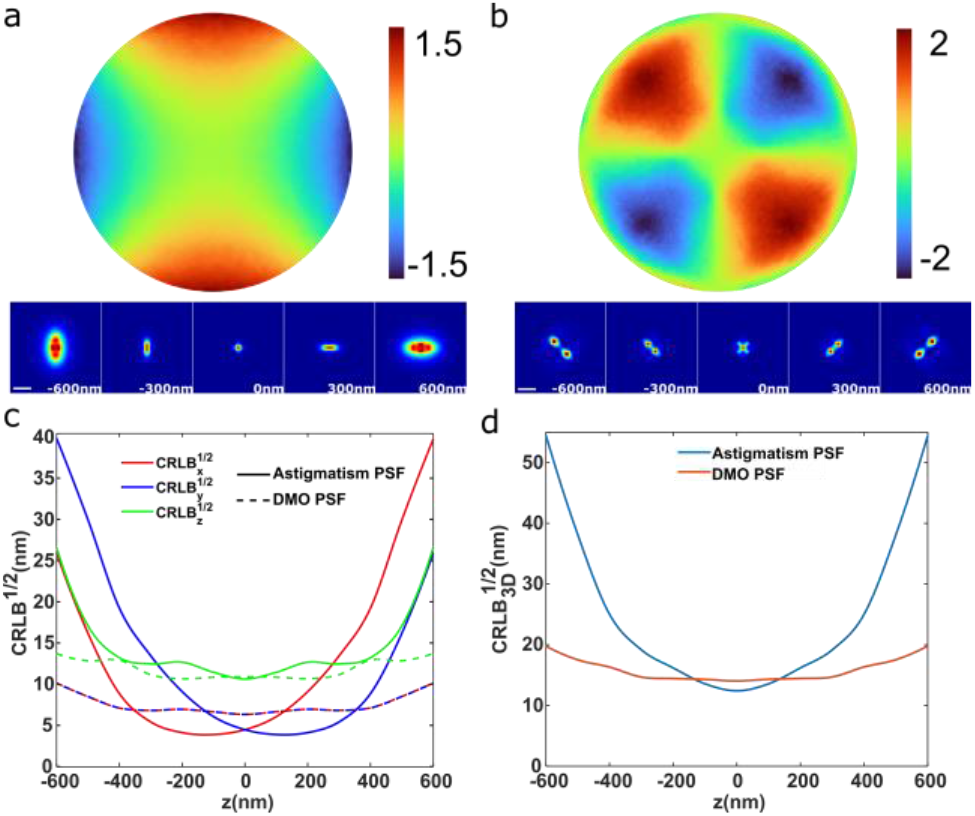
Comparison of astigmatism PSF and DMO Saddle-point PSF. (a) Astigmatism PSF (91 mλ rms). Projected pupil function on DM (top) and corresponding theoretical PSF (bottom). (b) DMO Saddle-point PSF as in (a). (c) The sqrt (CRLB_xyz_) of the astigmatism PSF and DMO Saddle-point PSF as a function of *z*. (d) The sqrt (CRLB_3D_) of the astigmatism PSF and DMO Saddle-point PSF as a function of *z*. All CRLB is calculated with a full vectorial PSF model [17] with the parameters: NA 1.5 for oil objective, 1.35 for silicone oil objective; pixel size 108 nm for x, y; wavelength 660 nm; refractive index (RI)1.518 for oil objective, 1.405 for silicone oil objective, assuming RI match for both cases; 2000 photons per emitter and a background of 20 photons per pixel. The same parameters are used throughout this work unless noted otherwise. Scale bars, 1μm.

To test the performance of the DMO Saddle-point PSF on the real biological sample, we imaged the nucleoporin Nup96 in U2OS cells using spline fitting method (**Supplementary Note 4**). Due to its stereotypical 3D structure, nuclear pore complex is widely used as a quantitative reference structure[18]. Here, we labeled the genome edited Nup96-SNAP cells with BG-AF647 fluorescent dyes. For comparison, we imaged Nup96 using both astigmatism and DMO Saddle-point 3D imaging methods (**Fig. 3 a** and **b**, **Visualization 1** and **2**). As shown in **Fig. 3 c, d, e**, and **f**, both methods could resolve the ring structure in *x-y* top view and bilayer structure in *x-z* cross section. However, in the zoomed image (**Fig. 3c** and **d**), the difference of the quality of the reconstructed images could be clearly observed. Close to the focus, both methods could resolve the individual Nup96 proteins in the symmetric unit of the NPC (white arrows in **Fig. 3c** and **d**). However, when it is slightly away from the focus (> 200 nm), the individual Nup96 protein can only be resolved by DMO Saddle-point PSF (red arrows in **Fig. 3 c** and **d**). This observation was further verified in the FRC analysis[19] (**Fig. 4**). For the regions close to focus (<140 nm), the FRC resolution is 31.1 nm and 31.6 nm for DMO Saddle-point PSF and astigmatism PSF, respectively. The FRC value for DMO Saddle-point PSF is almost constant between defocus regions (−500~−200 nm and 200~500 nm) and the near focal region, while it is more than twice as big (71.4 nm) for astigmatism PSF. This result is well in agreement with the theoretical calculation as shown in **Fig. 2**. Due to the superior performance of DMO Saddle-point PSF compared to astigmatism PSF, it has a great potential to totally replace the widely used astigmatism PSF.

**Fig. 3.**
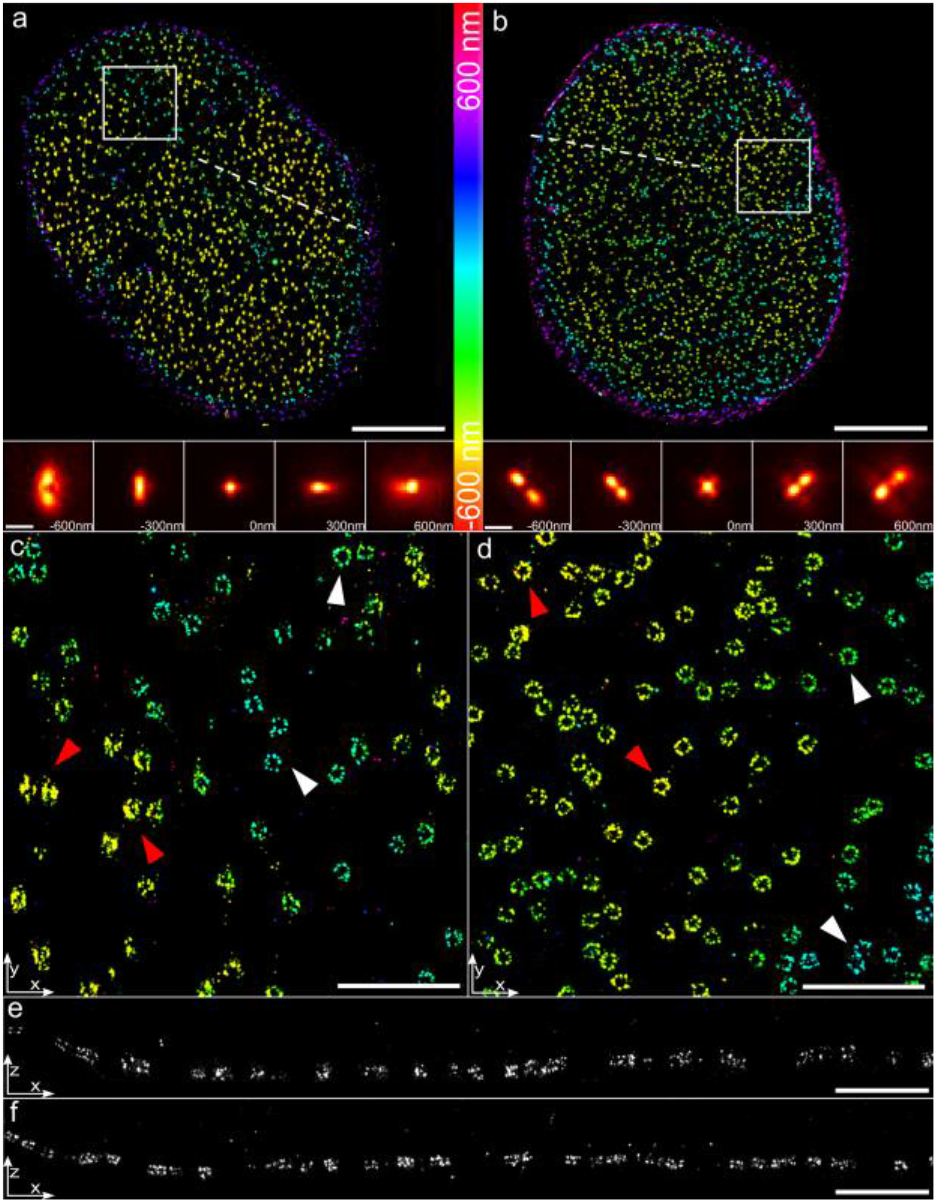
3D SMLM imaging of the Nup96-SNAP-AF647 using (a) astigmatism PSF and (b) DMO Saddle-point PSF with an NA 1.5 oil objective (UPLAPO100XOHR, Olympus). The bottom panels in (a) and (b) are the corresponding experimental PSFs. (c) and (d) are zoomed-in images of rectangles in (a) and (b), respectively. White and red arrows denote the near-focus and defocus NPCs, respectively. (e) and (f) are side view cross-sections the 500 nm thick lines denoted in (a) and (b), respectively. Scale bars, 5μm (top panels in a and b) and 1μm (bottom panels in a and b; c; d; e; f).

**Fig. 4.**
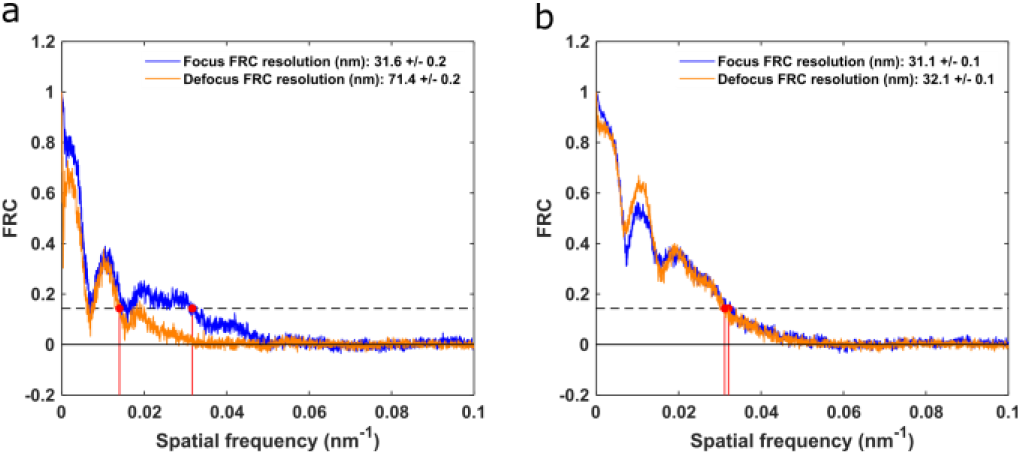
Fourier ring correlation (FRC) analysis of the super-resolution images within focus (+-140 nm) and defocus area (−500~−200 nm and 200~500 nm) imaged using (a) astigmatism PSF and (b) DMO Saddle-point PSF.

To further extend the depth of field of 3D super-resolution imaging, we then optimized a DMO Tetrapod PSF that has a 6-μm axial detection range which could image the entire nucleus without scanning. We first optimized a 6-μm theoretical Tetrapod PSF based on the Zernike polynomials and projected it on the DM to get a practical usage. Then a DMO Tetrapod PSF is optimized for further improvement. As shown in **Fig. S2 c** and **d**, the *CRLB*_3D, avg_ of the DMO Tetrapod PSF is improved by 8% compared to the projected one. To demonstrate the 3D resolving capability of the DMO Tetrapod PSF, we imaged the whole nucleus of Nup96 in the U2OS cells (**Fig. 5**, **Visualization 3**). To account for the RI mismatch, we used an NA 1.35 silicone oil objective (UPLSAPO100XS, Olympus) and embedded the sample in RI matched buffer using 2,2’-thiodiethanol. Since the relatively large size of the DMO Tetrapod PSF, the lateral overlap of the emitter PSFs is commonly observed. Here, we used the *R*_avg_(*PSF*) term (*α* = 30) to control the size of the optimized PSF. The distance between the two main lobes was reduced by ~20% compared to that of PSFs optimized without using *R*_avg_(*PSF*) term (*α* = 0) (**Fig. S3**). Deep-learning based algorithm, DECODE[15], was used to analyze the DMO Tetrapod PSF encoded single molecule data (**Supplementary Note 4**). As shown in **Fig. 5c** and **d**, the structure of the whole nuclear envelope could be nicely reconstructed. In the zoomed image as shown in **Fig. 5 e** and **f**, we were able to resolve the nuclear pores as rings both in the top and bottom of the nuclear envelope. The overall FRC resolution is 48.3 nm (**Fig. S4**).

**Fig. 5.**
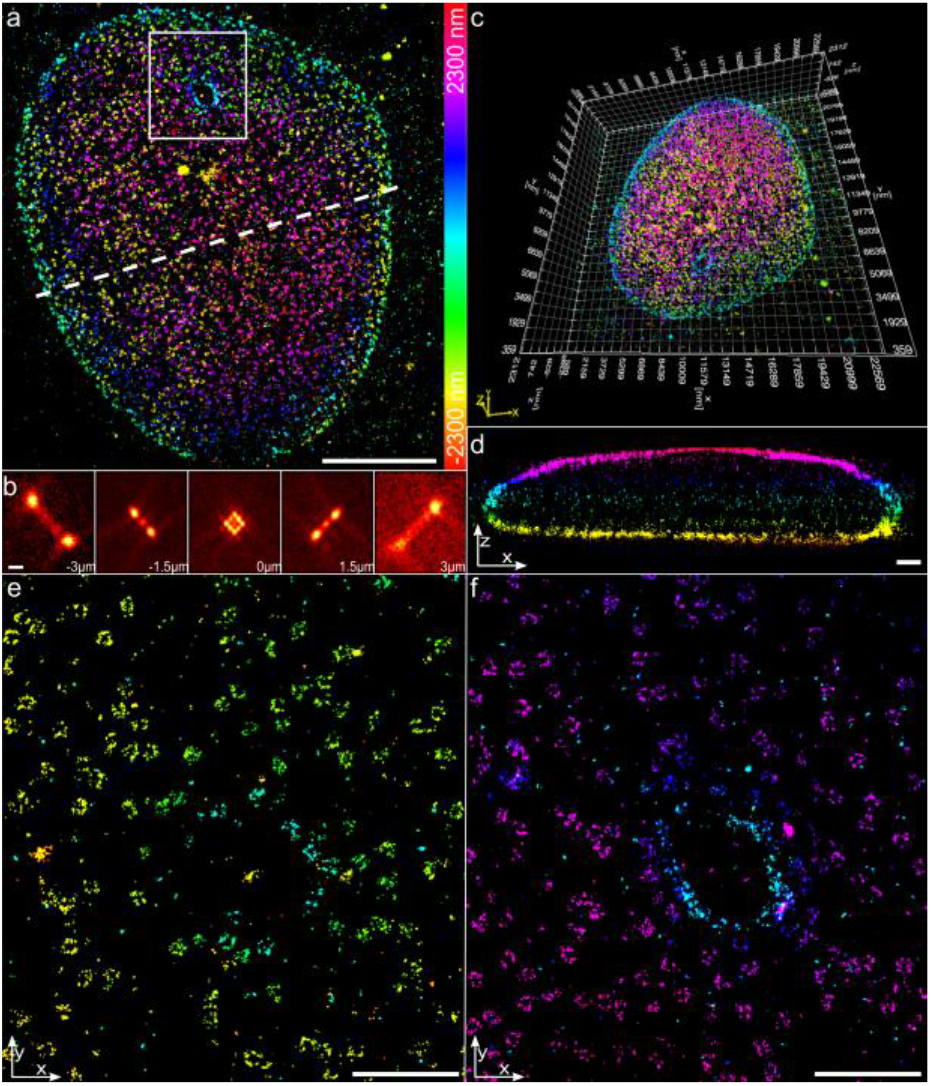
Whole-nucleus 3D super-resolution imaging of the Nup96-SNAP-AF647. (a) 3D super resolution images of the whole nucleus. (b) DMO Tetrapod experimental PSF. (c) Overview of a 3D reconstructed nucleus rendered by ViSP[20]. (d) Side view cross-section of the 2 μm thick line denoted in (a). (e) and (f) are zoomed views of the top surface and bottom surface of the nucleus, denoted by the rectangle in (a). Scale bars, 5μm (a) and 1μm (b; d; e; f).

In conclusion, we developed a DM specific PSF optimization method. In contrast to the previous method using Zernike polynomials as the solution space, we employed the influence functions of the DM as the solution space which could represent the behavior of the device best. To demonstrate the feasibility of our algorithm, we designed two DMO PSFs with different imaging depths: DMO Saddle point PSF and DMO Tetrapod PSF. We showed both in simulated and experimental data that DMO Saddle point PSF has almost twice better averaged CRLB than that of an astigmatism PSF, while maintaining a relatively constant resolution over the axial range being optimized (~1.2 μm). Furthermore, the DMO Tetrapod PSF could further improve the conventional “optimal” Tetrapod PSF by about 10% - 20 % on the DM and successfully super resolved the nuclear pores on the whole nuclear envelope. Our code is open source with detailed instructions in Ref. [21]. We believe that this work will greatly improve the widely used astigmatism PSF and generally improve the DM based PSF engineering.

## Data Availability

The code used in this work is available in Ref. [21].

## Supplemental Document

Supporting content in Supplement 1.

