## Supplemental information for "Deformable Mirror based Optimal PSF Engineering for 3D Super-resolution Imaging"

### Supplementary materials

| | |
|---|---|
| **Supplementary Fig. S1** | Layout of the custom-built microscope equipped with a deformable mirror |
| **Supplementary Fig. S2** | Comparison of CRLB of DM projected PSF and DMO PSF |
| **Supplementary Fig. S3** | Effect of the spatial confinement term to the shape of the PSF |
| **Supplementary Fig. S4** | Fourier ring correlation (FRC) analysis of the whole-nucleus 3D super-resolution image |
| **Supplementary Fig. S5** | Example steps for DM installation |
| **Supplementary Fig. S6** | Schematic of the imaging formation model of a fluorescent emitter |
| **Supplementary Note 1** | DM calibration and installation |
| **Supplementary Note 2** | Vectorial PSF model calculation |
| **Supplementary Note 3** | Derivation of CRLB and analytical gradients |
| **Supplementary Note 4** | Localization methods |
| **Supplementary Note 5** | Sample preparation |
| **Visualization 1** | Animated 3D view of the NPCs in **Fig. 3b** |
| **Visualization 2** | Animated 3D view of a representative single NPC in **Fig. 3b** |
| **Visualization 3** | Animated 3D view of the whole nucleus in **Fig. 5** |
| **Code 1** | Example and source code for DM based PSF optimization |

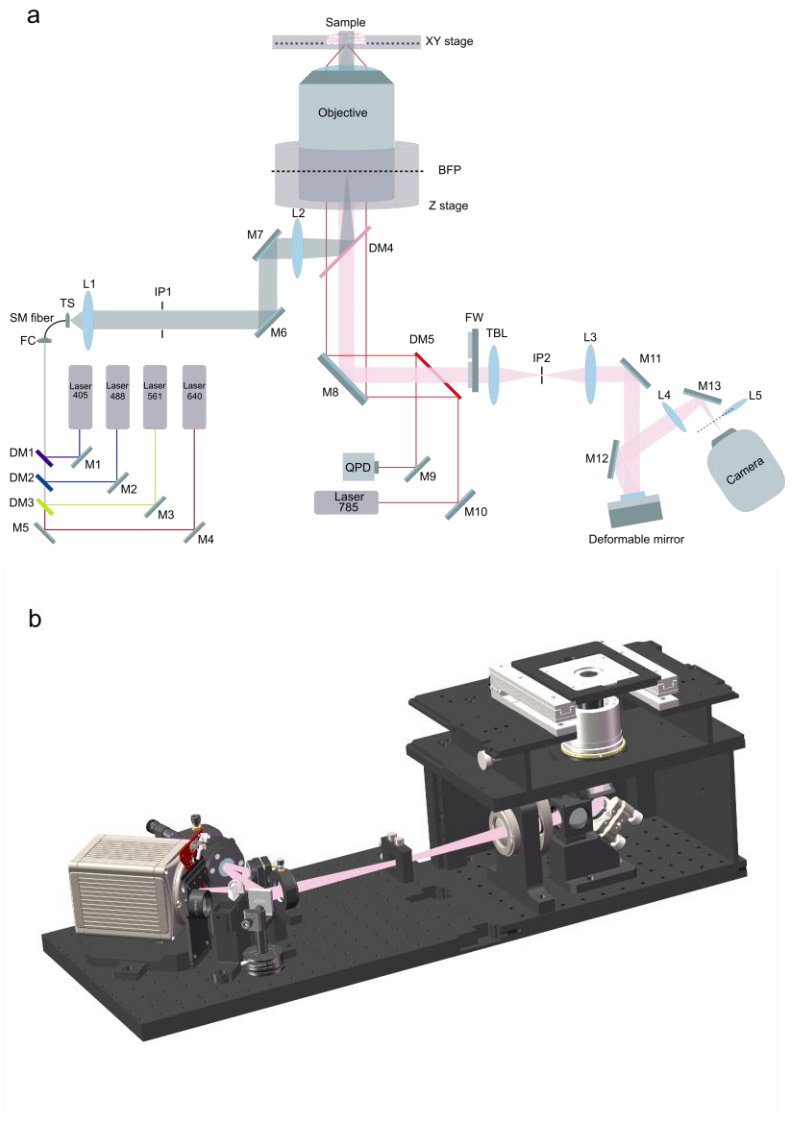


Supplementary Fig. S1. Layout of the custom-built microscope equipped with a deformable mirror. M: mirror, DM: dichroic mirror, L: lens, TS: translation stage, FC: fiber coupler, SM fiber: single-mode fiber, BFP: back focal plane, FW: filter-wheel, TBL: tube lens, IP: image plane, QPD: quadrant photodiode. (a) Beams from a 405 nm laser (IBEAM-SMART-405-S, 150 mW, TOPTICA Photonics), a 488 nm laser (IBEAM-SMART-488-S-HP, 200 mW, TOPTICA Photonics), a 561 nm laser (MGL-FN-561nm, 300mW, CNI) and a 640 nm laser (IBEAM-SMART-640-S-HP, 200mW, TOPTICA Photonics) are combined by a fiber coupler (PAF2-A4A, Thorlabs) and sent through a single-mode fiber (P3-405BPM-FC-2, Thorlabs). The fiber output position can be adjusted by the translation stage for different illumination angles. The illumination beam is filtered by a laser clean-up filter (ZET405/488/561/640xv2, Chroma) to remove fiber-induced fluorescence. A pair of lenses L1 (75 mm) and L2 (400 mm) with a slit (SP60, Owis) at IP1 are used for beam collimation and reshaping. The beam is then reflected by the main dichroic mirror (ZT405/488/561/640rpcxt-UF2, Chroma) before entering the objective for sample illumination. The emitted fluorescence is collected by a high NA objective (NA 1.35, UPLSAPO 100XS or NA 1.5, UPLAPO 100XOHR, Olympus) and imaged by the tube lens (TTL-180-A, Thorlabs) onto the IP2 confined by a slit (SP40, Owis). Two bandpass filters (NF03-405/488/561/635E-25 and FF01-676/37-25, Semrock) are used to separate the emitted fluorescence from the excitation laser. A 4-f system (L3, 125 mm, L4, 75 mm) with a deformable mirror (DM140A-35-P01, Boston Micromachines) placed in the Fourier plane, is set up for PSF engineering. Finally, images are acquired by an sCMOS camera (ORCA-Flash4.0 V3, HAMAMATSU) with pixel size of 108 nm in the sample. We typically chose to acquire 50,000-100,000 frames with a 15-ms exposure time. Besides, a closed-loop focus lock system is implemented, using the reflected signal of a 785 nm laser (IBEAM-SMART-785-S, 125 mW, TOPTICA Photonics) from the coverslip and its detection by a quadrant photodiode (SD197-23-21-041, Light Catcher), and achieved $\pm$10 nm focus stabilization over several hours.  (b) The rendered mechanical design of the microscope system using SolidWorks.

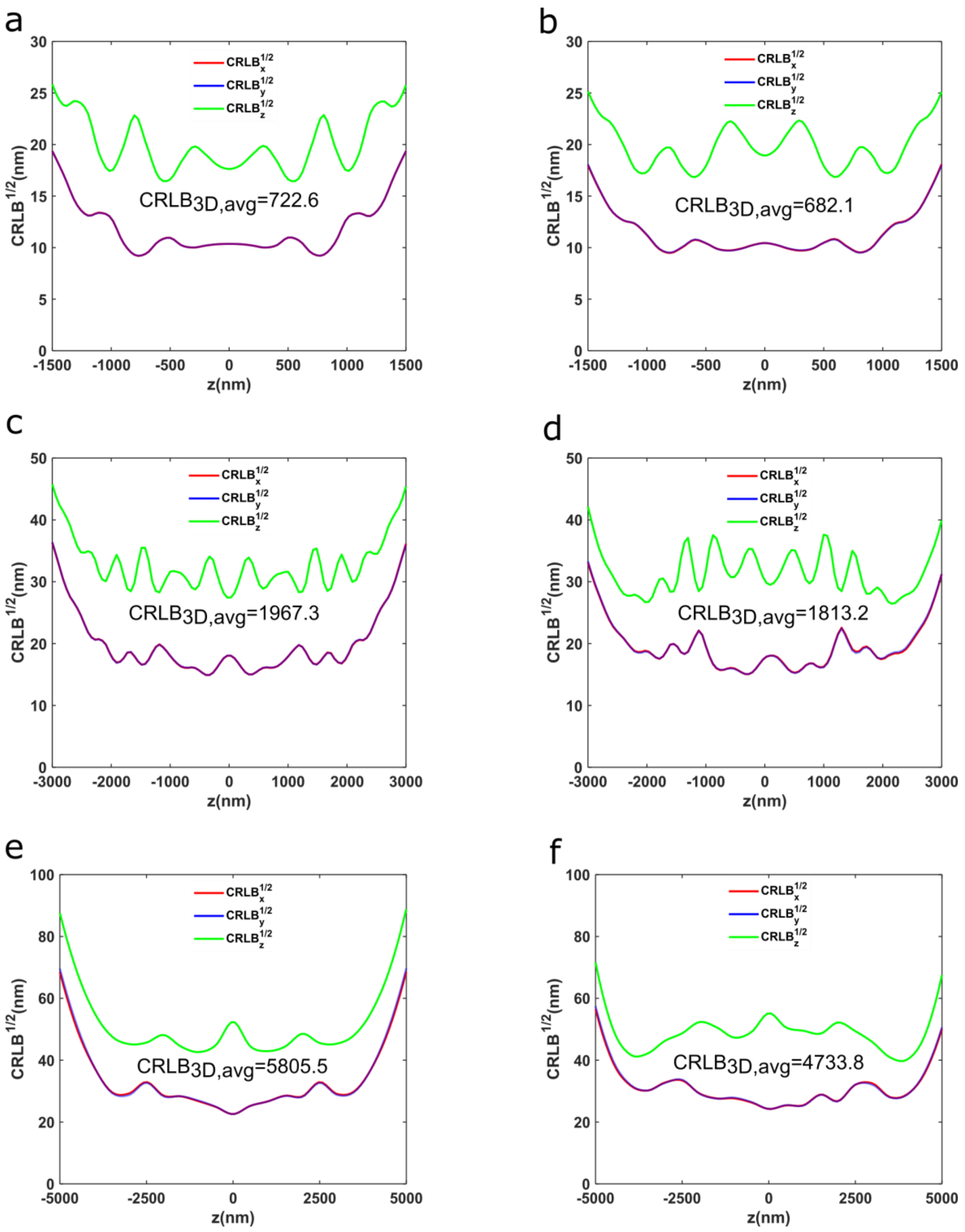


Supplementary Fig. S2. Comparison of CRLB of DM projected PSF and DMO PSF. The CRLB of the DM projected PSF using Zernike based optimization and DMO PSF optimized different axial ranges: (a) and (b) 3 µm; (c) and (d) 6 µm; (e) and (f) 10 µm. DM influence function-based optimizations (b, d, f) show better performance than Zernike-based optimizations (a, c, e).

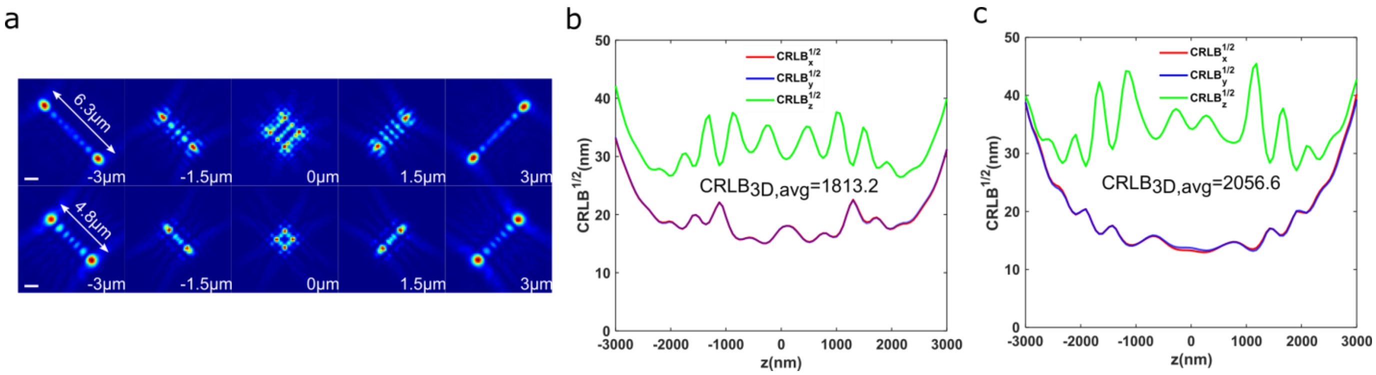


Supplementary Fig. S3. Effect of the spatial confinement term to the shape of the PSF. (a) Optimized PSFs in the 6 μm axial range with different spatial confinement term. Top row: DMO Tetrapod PSF optimized without using $R_{\text{avg}}(PSF)$ term ($\alpha = 0$). Bottom row: DMO Tetrapod PSF optimized using $R_{\text{avg}}(PSF)$ term ($\alpha = 30$). (b) and (c) are the sqrt ($\text{CRLB}_{\text{xyz}}$) for $\alpha = 0$ and $\alpha = 30$ as a function of z respectively. Scale bars, 1μm.

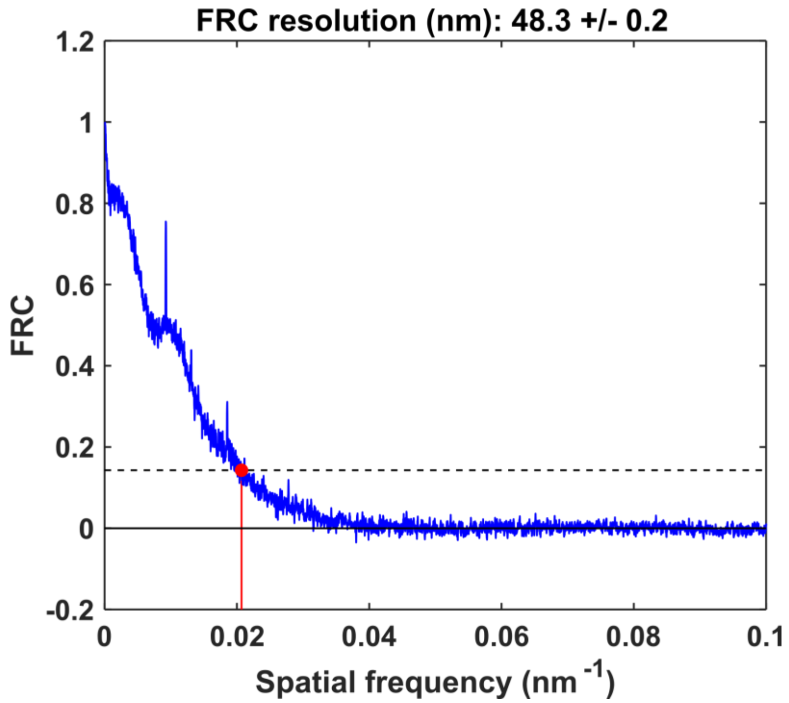

Supplementary Fig. S4. Fourier ring correlation (FRC) analysis of the whole-nucleus 3D super-resolution image in Fig. 5.

#### Supplementary Note 1: DM calibration and installation

##### *Principle of calibration*

To get accurate influence functions of each actuator of the DM (Boston Micromachines, DM140A-35-P01), we followed the method in Ref. [1]. First, we built a Twyman-Green interferometer, a variant of the Michelson interferometer, which is often used for optical testing. The phase information induced by the DM could be simply achieved by Fourier-based fringe analysis[2]. In this work, we assume that the responses of DM actuators are linear, and the inter-actuator coupling is neglected. Therefore, the output phase $\psi_{DM}$ can be expressed as a linear model:

$$\psi_{DM} = \sum \phi_m v_m \tag{S1}$$

where $\phi_m$ represents the influence function of the *m*-th actuator, $v_m$ is the control voltage applied to the *m*-th actuator. A sampled version of Eq. (S1) is:

$$\Psi_{DM} = \Phi \mathrm{V} \tag{S2}$$

Assuming the DM has $N_m$ actuators and $\Psi_{DM}$ is the discrete measured phase. The control signals V is a vector of size $N_m$ whereas $\Psi_{DM}$ is a vector of size $N_k$. The corresponding influence matrix $\Phi$ should be with a size $N_k \times N_m$, each column of $\Phi$ is a squeezed version of an influence function. After poking each actuator sequentially with different amplitudes and recording the interference images, we can calculate a series of output phases $[\Psi_1, \Psi_2 \dots \Psi_I]$ corresponding to their control signals $[\mathrm{V}_1, \mathrm{V}_2 \dots \mathrm{V}_I]$. Then the influence matrix $\Phi$ can be obtained by solving the simple least-squares problem:

$$\Phi = \underset{\Phi \in \mathbb{R}^{N_k \times N_m}}{argmin} \sum_I \|\Psi_i - \Phi \mathrm{V}_i\|^2 \tag{S3}$$

Once the influence matrix $\Phi$ is determined, we can easily know what control signals should be given to the DM to generate the desired wavefront shape through matrix calculation. We believe this can supply more accurate wavefront design than the wavefront generated by the calibrated Zernike modes as the Zernike calibration must hold the assumption:

$$\Psi_i \approx \mathbb{Z}\boldsymbol{\zeta} \tag{S4}$$

where $\mathbb{Z}$ is a matrix whose columns are Zernike polynomials sampled over the phase measurement grid. $\boldsymbol{\zeta}$ is a Zernike coefficients vector with size $N_{\boldsymbol{\zeta}}$ larger than $N_m$.

##### *Installation of DM*

The DM should be accurately installed in the Fourier plane of the microscope and conjugated to the objective pupil. The whole optical path of the system was optimized using Zemax so that the reflected membrane of the DM was placed on the Fourier plane. The incident light angle on the DM was designed to be 9.5 degree. In order to precisely put the center of the active membrane on the optical axis, the following steps were used:

1. Illuminate a fluorescent beads sample and check the fluorescence image to ensure that the optical path is normal.
2. Place the BFP lens (L5 in the Supplementary Fig. S1a) to the imaging path to image the objective pupil on the camera. Mark the center of the objective pupil (blue star, Supplementary Fig. S5a).
3. Switch off the excitation laser.
4. Use a white light source to illuminate the DM so that the DM can be clearly imaged on the camera with the BFP lens (Supplementary Fig. S5b).
5. Use the DM control software to apply a cross-pattern to the DM. Image the pattern on the camera and mark the center of the pattern (red star, Supplementary Fig. S5c).
6. Move the DM laterally until the cross-pattern's center overlaps with the center of the marked objective pupil (Supplementary Fig. S5d).

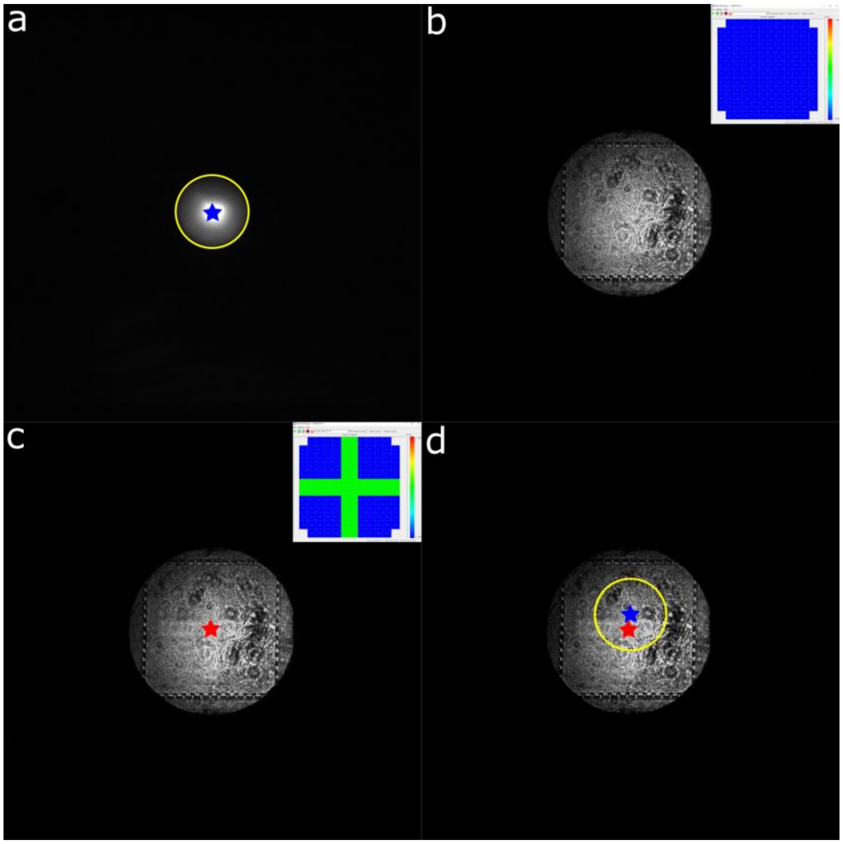


Supplementary Fig. S5. Example steps for DM installation. (a) Mark the center of the objective pupil. (b) Illuminate the DM with a bright light and image the DM on the camera. (c) Apply a cross-pattern to the DM. (d) Finetune the DM position until the pattern's center (red start) overlaps with the marked pupil center (blue star).

#### Supplementary Note 2: Vectorial PSF model calculation

To accurately describe the imaging formation process of the microscope with a high numerical aperture (NA) objective, a vectorial PSF model that takes account for refractive index mismatch between medium-cover slip interface and cover slip-immersion medium interface is used[3] . Since the fluorescent probes are normally attached to the molecules of interest flexibly and can rotate freely, we assumed an isotropic emitter PSF model. Taking the DM phase modulation into consideration, the PSF can be expressed as:

$$PSF \propto \sum_{p=x,y} \sum_{d=x,y,z} \left|E_{p,d}^{img}\right|^2, \ with \ E_{p,d}^{img} = \mathcal{F}_{2D}\left\{A(\rho,\varphi)e^{i\psi_{DM}(\rho,\varphi)}e^{i\psi_{pos}(\rho,\varphi;x_0,y_0,z_0,l)}E_{p,d}^{BFP}\right\} \qquad (S5)$$

where $E_{p,d}^{img}$ is the electric field with components $p = x, y$ in the image plane, and each dipole component $d = x, y, z$ in the sample contributes to components $p$ incoherently. $\mathcal{F}_{2D}$ denotes the 2D Fourier transform. $(\rho, \varphi)$ is the normalized polar coordinate in the back focal plane (BFP) whereas $\rho_{max}$ corresponds to the limiting aperture angle $NA/n_3$ and $\varphi$ corresponds to the azimuthal angle of the wave direction. $A(\rho, \varphi)$ is the aplanatic amplitude correction function $(1 - \rho^2 NA^2/{n_1}^2)^{-1/4}$. Here, NA is the numerical aperture. $n_1$, $n_2$ and $n_3$ are refractive index of sample medium, cover glass and immersion oil respectively. $\psi_{DM}$ is the extra phase induced by the DM as defined in Equation S2. In this work, we aim to find an optimal series of control voltages $v$ to engineer the PSF. $\psi_{pos}$ is the phase shift dependent on the emitter position:

$$\psi_{pos} = \frac{2\pi}{\lambda}\left(NAx_0\rho\cos\varphi + NAy_0\rho\sin\varphi + n_1 z_0\sqrt{1-(\frac{\rho NA}{n_1})^2} - n_3 l\sqrt{1-(\frac{\rho NA}{n_3})^2}\right) \qquad (S6)$$

where $\lambda$ is the wavelength, $(x_0, y_0, z_0)$ is the emitter position. $z_0$ represents the emitter's depth away from the cover glass. $l$ is the distance between the nominal focal plane and the cover glass (Supplementary Fig. S6). $E_{p,d}^{BFP}$ represents the polarization vectors in the BFP:

$$E_{x,d}^{BFP} = T_p\vec{P}_d\cos\varphi - T_s\vec{S}_d\sin\varphi \qquad (S7a)$$

$$E_{y,d}^{BFP} = T_p \vec{P}_d \sin\varphi + T_s \vec{S}_d \cos\varphi \quad \text{(S7b)}$$

$$with \left( \vec{P} = \begin{bmatrix} \cos\theta_1 \cos\varphi \\ \cos\theta_1 \sin\varphi \\ -\sin\theta_1 \end{bmatrix}, \vec{S} = \begin{bmatrix} -\sin\varphi \\ \cos\varphi \\ 0 \end{bmatrix} \right) \quad \text{(S7c)}$$

where $\vec{P}$ and $\vec{S}$ are the basis polarization vectors. $T_p$ and $T_s$ are the total Fresnel transmission coefficients for the p- and s-polarized light through multiple mediums: $T_p = T_{p,1-2} \times T_{p,2-3}, T_s = T_{s,1-2} \times T_{s,2-3}$. $\theta$ and $\varphi$ are the polar and azimuthal angles of the wave direction. Take the refractive index boundary between medium 1 and medium 2 as an example, the Fresnel transmission coefficients are given by:

$$T_{p,1-2} = \frac{2n_1 \cos\theta_1}{n_1 \cos\theta_2 + n_2 \cos\theta_1} \quad \text{(S8a)}$$

$$T_{s,1-2} = \frac{2n_1 \cos\theta_1}{n_1 \cos\theta_1 + n_2 \cos\theta_2} \quad \text{(S8b)}$$

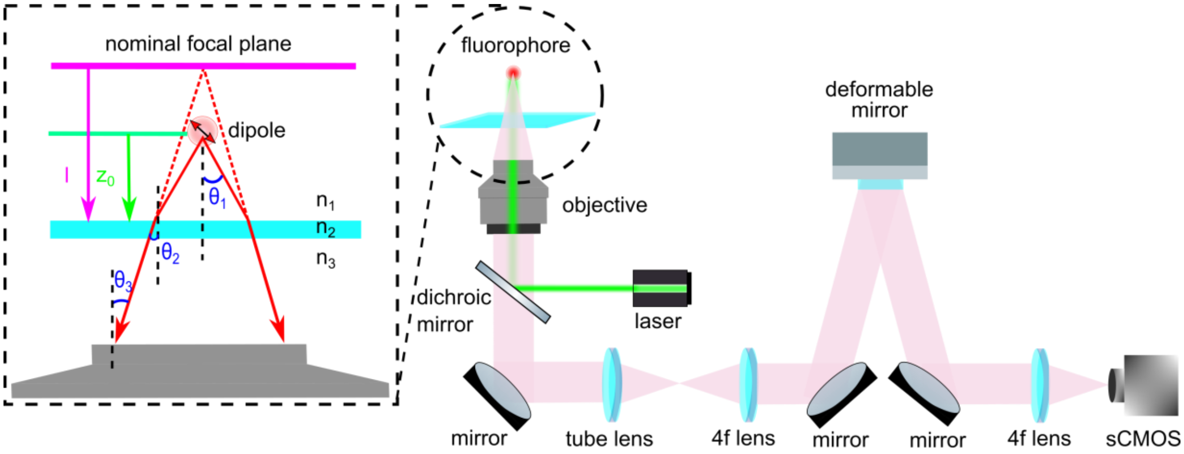


Supplementary Fig. S6. Schematic of the imaging formation model of the fluorescence microscope.

#### Supplementary Note 3: Derivation of CRLB and analytical gradients

To find a set of DM control signals $v$ that can engineer a PSF with the best localization performance, the objective function is defined as:

$$Loss = CRLB_{\mathrm{3D,\,avg}} + \alpha R_{\mathrm{avg}}(PSF) \quad \text{(S9a)}$$

$$= \frac{1}{N_z} \sum_{z \in \mathrm{Z}} \left( CRLB_{\hat{x},z} + CRLB_{\hat{y},z} + CRLB_{\hat{z},z} + \alpha R(PSF_z) \right) \quad \text{(S9b)}$$

$$= \frac{1}{N_z} \sum_{z \in \mathrm{Z}} \left( \left( \sum_{j=\hat{x},\hat{y},\hat{z}} [\mathrm{I}_z(\Theta)^{-1}]_{jj} \right) + \alpha R(PSF_z) \right) \quad \text{(S9c)}$$

The CRLB is calculated using the inverse of the Fisher information[4] matrix $\mathrm{I}_z(\Theta)$, which measures the amount of information that an observation (PSF) carries about the estimated parameters $\Theta$. Since we only optimize the precision of the position estimation, only the first three diagonal elements of $\mathrm{I}_z(\Theta)^{-1}$ are used (corresponding to $\hat{x}, \hat{y}, \hat{z}$ estimation). The Fisher information matrix is defined as:

$$\mathrm{I}(\Theta)_{ij} = \sum_k \frac{1}{u_k} \frac{\partial u_k}{\partial \Theta_i} \frac{\partial u_k}{\partial \Theta_j}, \quad \text{with } u_k = [\Theta_{photon} PSF + \Theta_{bg}]_k \quad \text{(S10)}$$

where $\Theta$ is a set of parameters being estimated. $\Theta_x, \Theta_y, \Theta_z, \Theta_{photon}$ and $\Theta_{bg}$ are corresponding to $x$, $y$, $z$, photon, and background, respectively. $\mathrm{I}(\Theta)$ is a $5 \times 5$ matrix. $u_k$ is the expected photons in pixel $k$ of the pixelated PSF model. In order to speed up the optimization process, the explicit gradient function of the loss function was derived. The gradient of the objective function with respect to the DM control signal is given by:

$$\frac{\partial Loss}{\partial v_m} = \frac{1}{N_z}\sum_{z\in Z}\left(\left(\sum_{j=\hat{x},\hat{y},\hat{z}}\left[-\mathrm{I}_z(\Theta)^{-1}\frac{\partial \mathrm{I}_z(\Theta)}{\partial v_m}\mathrm{I}_z(\Theta)^{-1}\right]_{jj}\right) + \alpha R\left(\frac{\partial PSF_z}{\partial v_m}\right)\right) \tag{S11}$$

where $v_m$ is the control voltage applied to the *m*-th DM actuator. Following the chain rule, the gradient of Fisher matrix elements can be expressed as:

$$\frac{\partial \mathrm{I}(\Theta)_{ij}}{\partial v_m} = \sum_k \left[\frac{\partial u_k}{\partial \Theta_i}\cdot\frac{\partial u_k}{\partial \Theta_j}\cdot\frac{-1}{{u_k}^2}\cdot\frac{\partial u_k}{\partial v_m} + \frac{1}{u_k}\cdot\frac{\partial u_k}{\partial \Theta_j}\cdot\frac{\partial^2 u_k}{\partial \Theta_i \partial v_m} + \frac{1}{u_k}\cdot\frac{\partial u_k}{\partial \Theta_i}\cdot\frac{\partial^2 u_k}{\partial \Theta_j \partial v_m}\right] \tag{S12}$$

The first derivatives of the PSF model $u_k$ with respect to the parameters $\Theta$ are given by:

$$\frac{\partial u_k}{\partial \Theta_{x,y,z}} = \frac{\Theta_{photon}}{C}\sum_{p=x,y}\sum_{d=x,y,z} Re\{E_{p,d}^{img^*}\frac{\partial E_{p,d}^{img}}{\partial \Theta_{x,y,z}}\} \tag{S13a}$$

$$\frac{\partial u_k}{\partial v_m} = \frac{\Theta_{photon}}{C}\sum_{p=x,y}\sum_{d=x,y,z} Re\{E_{p,d}^{img^*}\frac{\partial E_{p,d}^{img}}{\partial v_m}\} \tag{S13b}$$

$$\frac{\partial u_k}{\partial \Theta_{photon}} = PSF(k) \tag{S13c}$$

$$\frac{\partial u_k}{\partial \Theta_{bg}} = 1 \tag{S13d}$$

$$\frac{\partial E_{p,d}^{img}}{\partial \Theta_{x,y,z}} = \mathscr{F}_{2D}\{i\vec{k}_{x,y,z}A(\rho,\varphi)e^{i\psi_{DM}(\rho,\varphi)}e^{i\psi_{pos}(\rho,\varphi;\Theta)}E_{p,d}^{BFP}\} \tag{S13e}$$

$$\frac{\partial E_{p,d}^{img}}{\partial v_m} = \mathscr{F}_{2D}\{i\phi_m A(\rho,\varphi)e^{i\psi_{DM}(\rho,\varphi)}e^{i\psi_{pos}(\rho,\varphi;\Theta)}E_{p,d}^{BFP}\} \tag{S13f}$$

where $C$ is a normalization factor. $PSF(k)$ is the value of pixel $k$ of the normalized PSF model. $\vec{k}$ is the wavevector ($\vec{k} = 2\pi/\lambda(NA\rho\cos\varphi, NA\rho\sin\varphi, \sqrt{n^2-\rho^2 NA^2})$). $\phi_m$ is the influence function of the *m*-th actuator, and the second derivatives of the model are as follows:

$$\frac{\partial^2 u_k}{\partial \Theta_{x,y,z}\partial v_m} = \frac{\Theta_{photon}}{C}\sum_{p=x,y}\sum_{d=x,y,z} Re\{\frac{\partial E_{p,d}^{img^*}}{\partial v_m}\frac{\partial E_{p,d}^{img}}{\partial \Theta_{x,y,z}} + E_{p,d}^{img^*}\frac{\partial^2 E_{p,d}^{img}}{\partial \Theta_{x,y,z}\partial v_m}\} \tag{S14a}$$

$$\frac{\partial^2 u_k}{\partial \Theta_{photon}\partial v_m} = \frac{1}{C}\sum_{p=x,y}\sum_{d=x,y,z} Re\{E_{p,d}^{img^*}\frac{\partial E_{p,d}^{img}}{\partial v_m}\} \tag{S14b}$$

$$\frac{\partial^2 u_k}{\partial \Theta_{bg}\partial v_m} = 0 \tag{S14c}$$

$$\frac{\partial^2 E_{p,d}^{img}}{\partial \Theta_{x,y,z}\partial v_m} = \mathscr{F}_{2D}\{-\phi_m\vec{k}_{x,y,z}A(\rho,\varphi)e^{i\psi_{DM}(\rho,\varphi)}e^{i\psi_{pos}(\rho,\varphi;\Theta)}E_{p,d}^{BFP}\} \tag{S14d}$$

With the above analytical gradient of the objective function, there are a lot of algorithms that can be implemented to solve this optimization problem [5]. In this work we used the interior-point method of the built-in *fmincon* function in Matlab as we found it is convenient and robust to find a minimum. It should be noted that multiple starting parameters are needed to find the final optimal DM control signals.

#### Supplementary Note 4: Localization methods

##### *Maximum likelihood estimation of single molecule data*

For DMO Saddle-point PSF imaging, the data was analyzed with the cubic spline fitting method[6]. With the optimized voltages given to the DM, we acquired an experimental 3D PSF by averaging the bead stack images from different fields of view. Then the cubic spline functions were used to interpolate this 3D experimental PSF:

$$f_{i,j,k}(x,y,z) = \sum_{m=0}^{3}\sum_{n=0}^{3}\sum_{p=0}^{3} a_{i,j,k,m,n,p}\left(\frac{x-x_i}{\Delta x}\right)^m\left(\frac{y-y_j}{\Delta y}\right)^n\left(\frac{z-z_k}{\Delta z}\right)^p \tag{S15}$$

where $\Delta x$, $\Delta y$ are the xy pixel sizes, $\Delta z$ is the axial step size of the PSF stack, $a_{i,j,k,m,n,p}$ are the spline coefficients and $x_i, y_j, z_k$ are the start positions for each voxel $(i,j,k)$. After building the spline PSF model, maximum likelihood estimation with Poisson statistics was used to localize single molecules with the objective function given by:

$$\chi_{mle}^2 = 2\left(\sum_k (\mu_k - M_k) - \sum_{k,M_k>0} M_k \ln\left(\frac{\mu_k}{M_k}\right)\right) \quad (S16)$$

where $\mu_k$ and $M_k$ are the expected photon number and measured photon number in the kth pixel, respectively. We used a modified Levenberg-Marquardt (L-M) algorithm to minimize $\chi_{mle}^2$ for the parameter estimation. For DMO Saddle-point PSF imaging, molecules with $xy$ localization precision larger than 30 nm were removed. Molecules close than 35 nm within 3 adjacent frames were grouped as one molecule.

##### *Deep learning method used for large-DOV localization*

For DMO Tetrapod PSF imaging, deep-learning based method, DECODE[7], was used to analyze the easily overlapped and low-SNR Tetrapod single molecule data. The network was trained with an online data generator using an experimental spline PSF model. The input of the network is a set of three consecutive frames that mimic the temporal dynamics of single molecules. We set the noise model of simulated images as the sCMOS camera data, with parameters provided by the manufacture. The network outputs uncertainties for each molecule localization, which was used to filter bad localizations. The training was performed on simulated $128 \times 128$ images with a batch size of 10 and stopped until 30,000 iterations. The other training details and inferring procedures are the same in the Ref. [7]. For DMO Tetrapod PSF imaging, molecules with $xy$ localization precision larger than 50 nm were rejected. Molecules close than 100 nm within 3 adjacent frames were grouped as one molecule. Redundant cross-correlation algorithm[8] was used for drift correction (10 time windows were used in this work). For super-solution image rendering, we used SMAP[9] to produce all static super-resolution images, and the ViSP[10] software to render 3D movies.

#### Supplementary Note 5: Sample preparation

##### *Cell culture*

U2OS cells (Nup96-SNAP no. 300444, Cell Line Services) were grown in DMEM (catalog no. 10569, Gibco) containing 10% (v/v) fetal bovine serum (catalog no. 10270-106, Gibco), 1× MEM NEAA (catalog no. 11140-050, Gibco),100 U/ml penicillin and 100 μg/ml streptomycin (catalog no. 15140-122, Gibco). Cells were cultured in a humidified atmosphere with 5% $CO_2$ at 37 °C and passaged every two or three days. Prior to cell plating, high-precision 25-mm-round glass coverslips (no. 1.5H, catalog no. CG15XH, Thorlabs) were cleaned by sequentially sonicating in 1 M potassium hydroxide (KOH), Milli-Q water and ethanol, and finally irradiated under ultraviolet light for 30 min. For super-resolution imaging, U2OS cells were cultured on the clean coverslips for 2 d with a confluency of 50-70%.

##### *SNAP-tag labeling of Nup96*

To label Nup96, U2OS-Nup96-SNAP cells were prepared as previously reported[11]. Briefly, cells were prefixed in 2.4% paraformaldehyde (PFA) for 30 s, permeabilized in 0.4% Triton X-100 for 3 min and subsequently fixed in 2.4% PFA for 30 min. Then, cells were quenched in 0.1 M $NH_4Cl$ for 5 min and washed twice with PBS. To decrease unspecific binding, cells were blocked for 30 min with Image-iT FX Signal Enhancer (catalog no. I36933, Invitrogen). For labeling, cells were incubated in dye solution (1 μM BG-AF647 (catalog no. S9136S, New England Biolabs), 1 mM DTT and 0.5% bovine serum albumin (BSA) in PBS) for 1 h, and washed 3 times in PBS for 5 min each to remove excess dyes. Lastly, cells were postfixed with 4% PFA for 10 min, washed with PBS 3 times and stored at 4 ℃ until imaged.
